# Temperature restriction in entomopathogenic bacteria

**DOI:** 10.1101/2020.06.02.129163

**Authors:** Alexia Hapeshi, Joseph R.J. Healey, Geraldine Mulley, Nicholas R. Waterfield

**Affiliations:** Microbiology and Infection Unit, Warwick Medical School, The University of Warwick, Gibbet Hill Road, Coventry, CV4 7AL, UK; School of Biological Sciences, University of Reading, Whiteknights, Reading RG6 6AJ, UK

**Keywords:** temperature, adaptation, enterobacteria, *Photorhabdus*, symbionts, pathogens

## Abstract

Temperature plays an important role in bacteria-host interactions and can be a determining factor for host switching. In this study we sought to investigate the reasons behind growth temperature restriction in the entomopathogenic enterobacterium *Photorhabdus. Photorhabdus* has a complex dual symbiotic and pathogenic life cycle. The genus consists of 19 species but only one subgroup, previously all classed together as *P. asymbiotica*, have been shown to cause human disease. These clinical isolates necessarily need to be able to grow at 37 °C, whilst the remaining species are largely restricted to growth temperatures below 34 °C and are therefore unable to infect mammalian hosts. Here, we have isolated spontaneous mutant lines of *P. laumondii* DJC that were able to grow up to 36 °C-37 °C. Following whole genome sequencing of 29 of these mutants we identified a single gene, encoding a protein with a RecG-like helicase domain, that for the majority of isolates contained single nucleotide polymorphisms. Importantly, provision of the wild-type allele of this gene in *trans* restored the temperature restriction, confirming the mutations are recessive, and the dominant effect of the protein product of this gene. The gene appears to be part of a short three cistron operon, which we have termed the Temperature Restricting Locus (TRL). Transcription reporter strains revealed that this operon is induced upon the switch from 30 °C to 37 °C, leading to replication arrest of the bacteria. TRL is absent from all of the human pathogenic species so far examined, although its presence is not uniform in different strains of the *P. luminescens* subgroup. In a wider context, the presence of this gene is not limited to *Photorhabdus*, being found in phylogenetically diverse proteobacteria. We therefore suggest that this system may play a more fundamental role in temperature restriction in diverse species, relating to as yet cryptic aspects of their ecological niches and life cycle requirements.

## 1 Introduction

*Photorhabdus* is a Gram-negative bacterium that forms a symbiotic relationship with *Heterorhabditis* nematodes. The *Heterorhabditis*-*Photorhabdus* complex is entomopathogenic with the nematodes infecting insects and releasing the bacteria by regurgitation (Ciche and Ensign 2003). *Photorhabdus* then sets up a lethal septicaemia, producing an array of toxins and degradative enzymes that combat the immune system and kill the insect. They also elaborate a potent cocktail of antimicrobial compounds to eliminate competition by other microbes in the carcass. The bacteria reproduce using nutrients derived from the insect tissues, reaching very high numbers. The nematodes feed on this bacterial biomass, going through several rounds of hermaphroditic reproduction, before eventually re-establishing the symbiosis with the bacteria (reviewed in (Waterfield, Ciche, and Clarke 2009)). The *Photorhabdus* genus, which belongs to the class of γ-proteobacteria, was thought to consist of three species *P. asymbiotica, P. luminescens* and *P. temperata*. Recently, however, it has been proposed that several of the subspecies be elevated to species level resulting in a total of 19 species (Machado et al. 2018). Nevertheless, for the purposes of this study we will refer to these as belonging to the *P. asymbiotica, P. luminescens* and *P. temperata* subgroups. Thus, so far, only species belonging to the *P. asymbiotica* subgroup (that can grow up to 37 °C or above) have been identified as the causative agent of human infections (Gerrard and Stevens 2017). The only exception to this was a strain of *P. luminescens*, identified using 16S sequencing, which was associated with cutaneous lesions and bacteremia in a hypothermic infant with a body temperature upon admission of 33.6 °C (Dutta et al. 2018). No other members of the subgroups *P. luminescens* and *P. temperata*, the majority of which are largely restricted to growth temperatures below 37 °C (Fischer-Le Saux et al. 1999; Tailliez et al. 2010; Machado et al. 2018), have been reported to cause human disease. The exact reasons behind this discrepancy are unknown.

The establishment and maintenance of symbiosis is also a very important aspect of the *Photorhabdus* life cycle. There is considerable partner preference in the association between *Photorhabdus* bacteria and the *Heterhorhabditis* nematodes, although this is not strictly species-specific (Maher et al. 2017; Kazimierczak et al. 2017). Nonetheless, there is evidence for the co-evolution of the two partners (Maneesakorn et al. 2011). It may therefore be of relevance, that the optimum temperature for the nematodes is also relatively low. For example, in our hands the *P. laumondii* DJC *Heterorhabditid* nematode partner dies in culture above 34 °C (unpublished data).

*P. laumondii* sbsp. *laumondii* DJC is a laboratory strain of *Photorhabdus* which is normally unable to grow at temperatures above 35 °C on solid LB media. In this study we have isolated mutants of *P. laumondii* DJC with abilities to grow between 36-37 °C.

## 2 Results

### 2.1 Isolation and characterisation of temperature tolerant *P. laumondii* DJC mutants

Clones of *P. laumondii* DJC were isolated with the ability to grow on LB agar at 36 °C. This was following plating of a saturated overnight culture and incubation of the plates at the higher temperature. The process was repeated two more times and colonies were isolated from all three experiments originating from four independent lineages in total. Isolates were re-streaked 5 times at 36 °C to select for stably growing clones. Overall, the average frequency with which stable clones were isolated was 2.6 ± 2.5 ×10^−6^. Thirty of those were then analysed by whole genome sequencing to identify any mutations that may give rise to the phenotype. Sequencing was performed using the MiSeq platform and good quality reads were obtained for 29 of the 30 clones. The WT strain in the lab was also sequenced to account for variations between isolates in different laboratories. Reads from the temperature tolerant clones were then mapped against the reference to identify single nucleotide polymorphisms (SNPs).

Of the 29 isolates successfully sequenced, 19 had a SNP in an uncharacterised gene *PluDJC_RS01875* (Figure 1A and Table 1), and an additional isolate (S15) had an IS30-family transposase insertion near the 3’ end (chromosomal position 393017). This gene appears in the automated annotation as “*recG”*, however it should be noted that a full-length orthologue of the bona fide *recG* is encoded elsewhere in the genome. RecG is a 693 amino acid protein whose homologue in other bacteria has been well studied and is known to be important for DNA repair and recombination (Ishioka, Iwasaki, and Shinagawa 1997; Rudolph et al. 2010). As *PluDJC_RS01875* is involved in the temperature restriction phenotype, and to distinguish it from the *recG* misnomer, we subsequently named it *trlG* (***t***emperature ***r***estriction ***l***ocus **G**).

**Table 1.**
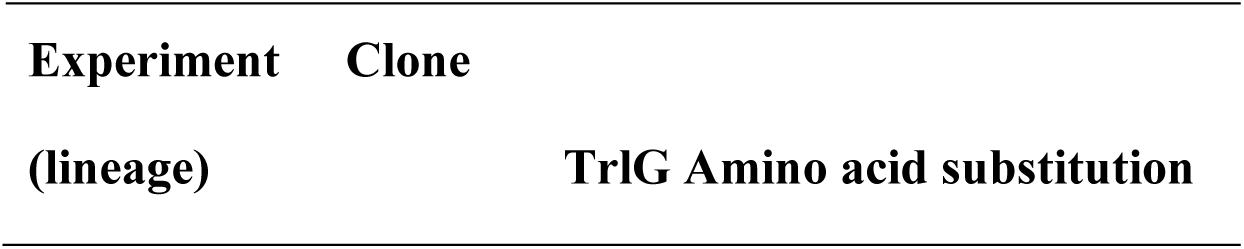
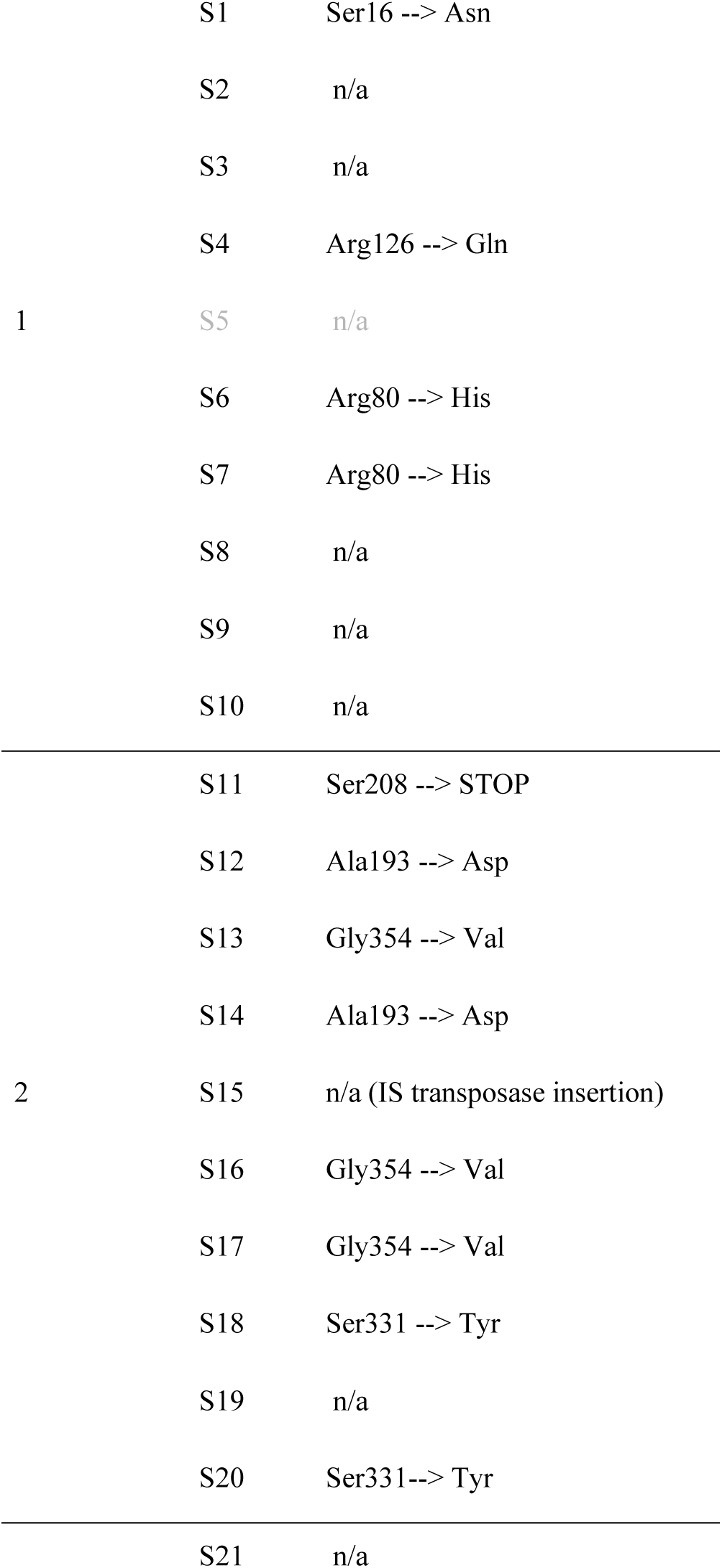
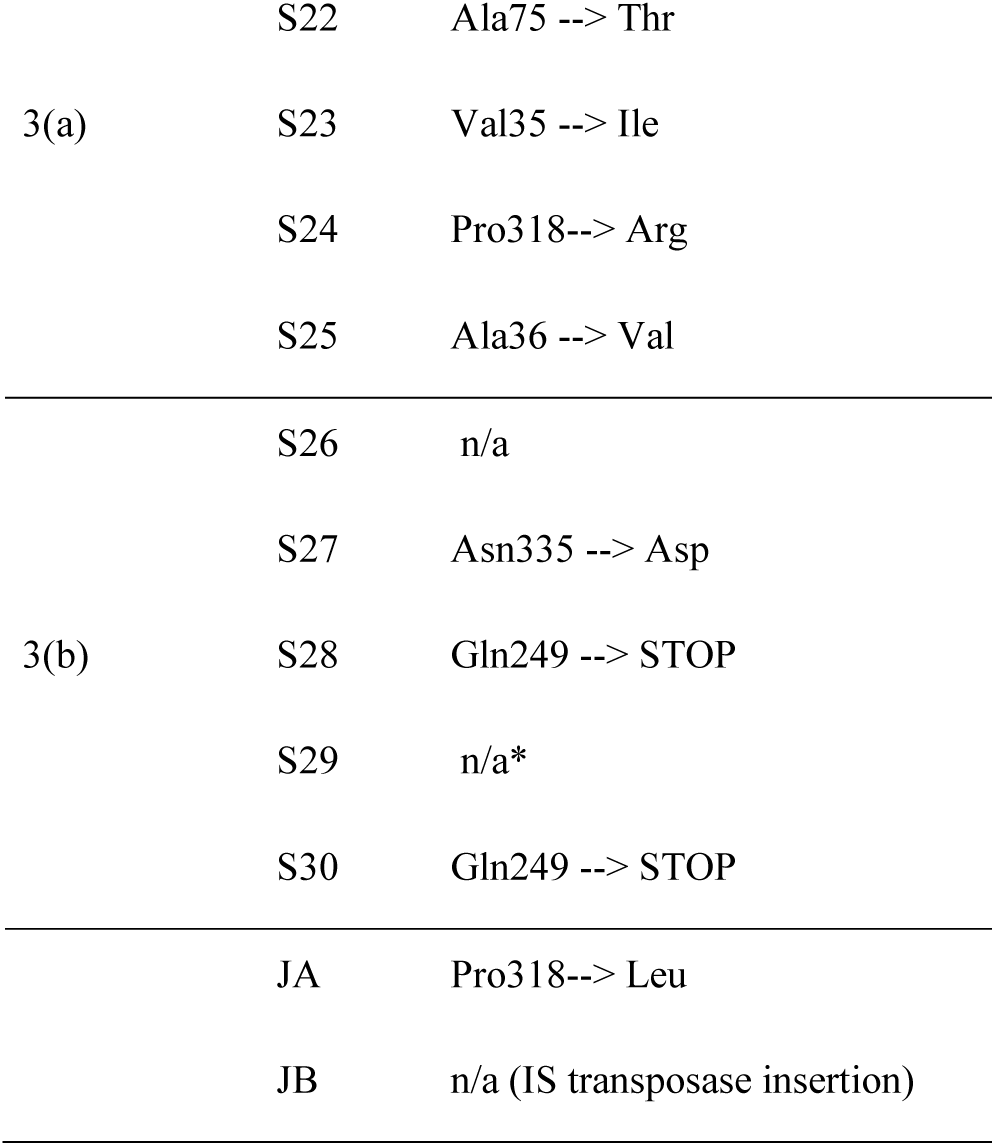
TrlG amino acid substitutions that result from the SNPs identified in the sequenced isolates, as well as isolates JA and JB. The isolates are separated based on the experiment and thus the lineage they originate from. Note that in experiment 3 two independent starting cultures were used. *Inspection of the aligned sequencing reads for clone S29 revealed possible SNPs or small insertions/deletions at the 3’ of *trlG*, however it was not possible to confirm that with Sanger sequencing of the amplified region.

**Figure 1.**
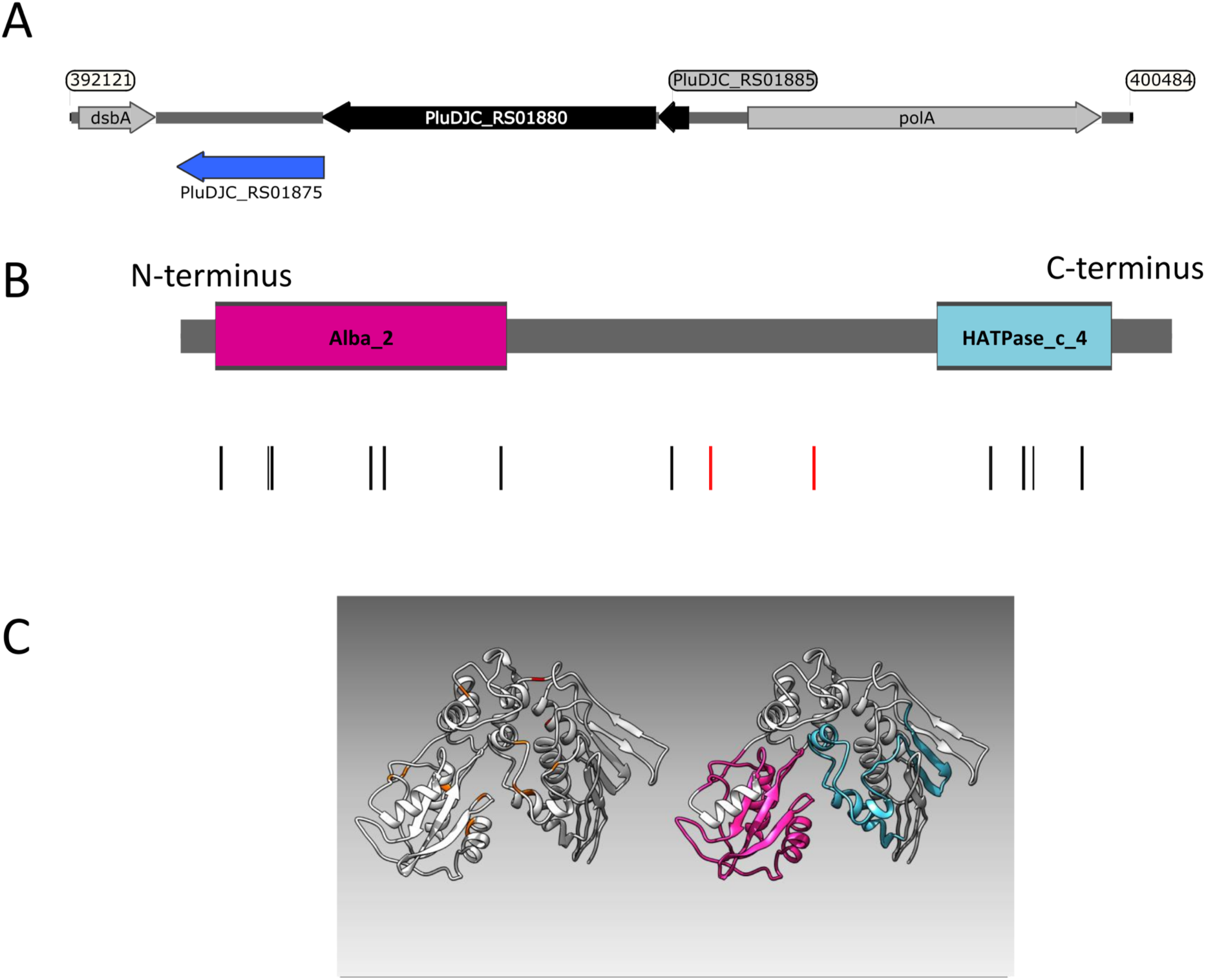
A) Genetic organisation of the Temperature restriction locus. B) Domain architecture of TrlG and the presence of substitution mutations in the temperature tolerant clones. Vertical lines indicate the positions of the mutations; red depicts stop mutations. The coloured boxes show the domains identified by Pfam; Magenta: Alba_2 (Pfam domain PF04326, at amino acid positions 14-128) and light blue: HATPase_c_4 (Pfam domain PF13749, at amino acid positions 298-365). C) **(Left)** I-TASSER simulation of TrlG with the distribution of mutations mapped. Orange indicates SNP positions, whilst red depicts stop mutations. **(Right)** The same model as in (Left) with homology domains mapped in colour. Images prepared with UCSF Chimera.

Repetition of these experiments aimed at isolating temperature tolerant clones by a collaborator in an independent laboratory (Dr G. Mulley), gave rise to two additional tolerant isolates, named JA and JB. These were specifically tested for the presence of mutations in *trlG* by sequencing of a PCR amplicon. Clone JA was found to contain a SNP, while in clone JB an IS66 transposable element had inserted into the 3’ end of *trlG* (chromosomal position 393014). Homology and structural models of the TrlG protein suggest it comprises two distinct functional domains connected by a flexible loop region. Most of the mutant SNPs were located in either of the two domains. Additionally, three mutants exhibited premature stop codon mutations in the linker region (Figure 1B). Based on a 3D structure prediction of the protein, we can observe that four of the SNPs would result in substitutions in alpha helices potentially disrupting the structure (Figure 1C). The remaining isolates that did not have a mutation in *trlG* were actually unable to resume growth at 36 °C after being frozen (Figure 2A and Supplementary Figure 1). It should be noted that the tolerant isolates are less pigmented and produce less bioluminescence at the higher temperature (Figure 2A and Supplementary Figure 2). We then investigated the hypothesis that the operon might affect growth or survival at colder temperatures by incubating the bacteria at lower temperatures before shifting back to 28 °C. However, the mutants in this case behaved similarly to the WT bacteria (data not shown).

**Figure 2.**
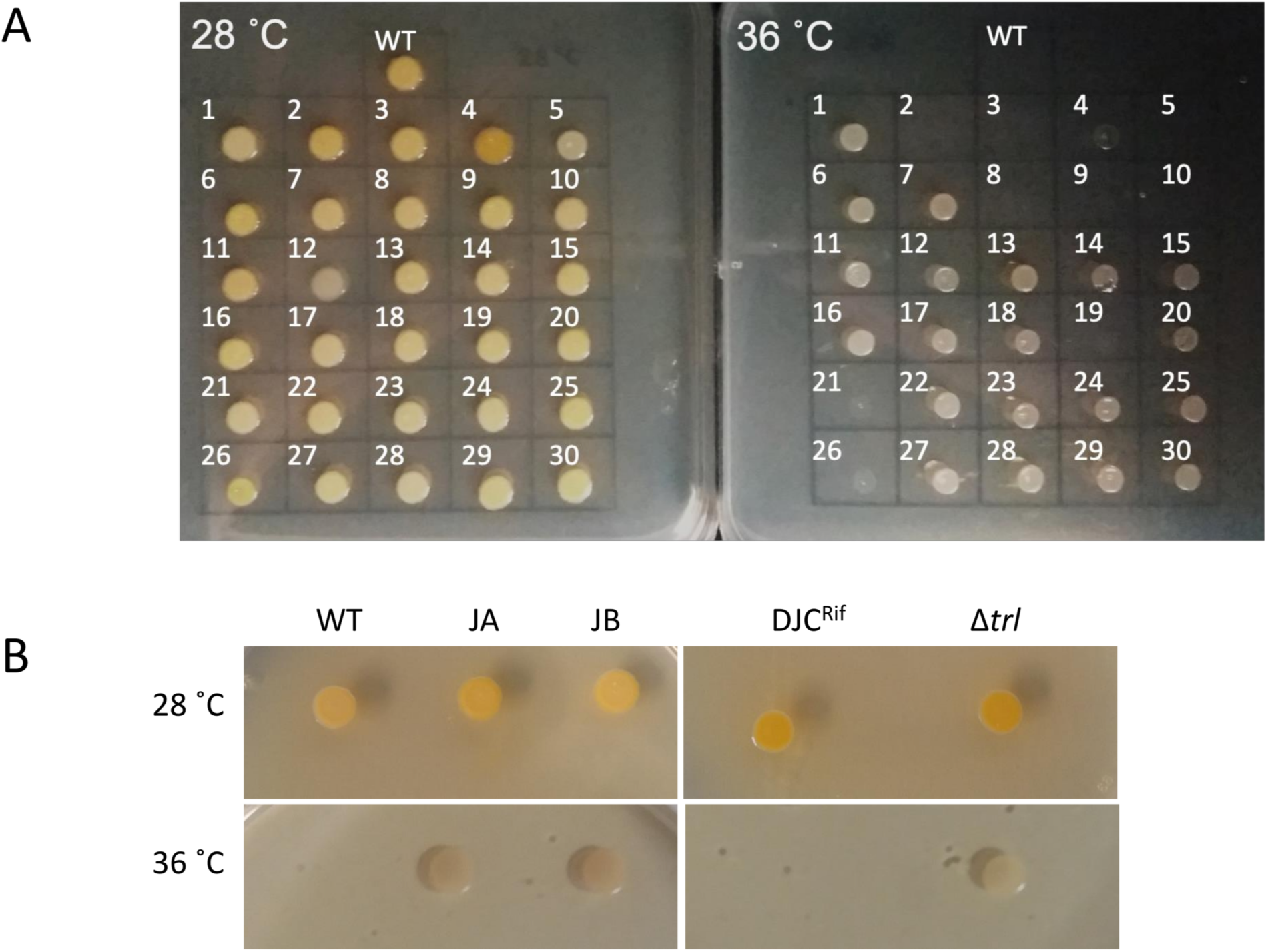
A) Growth of the 30 sequenced tolerant clones and *P. laumondii* DJC (WT) on LB agar at 28 °C and 36 °C. B) Growth of the additional two clones isolated independently (JA and JB) compared to *P. laumondii* DJC (WT) as well as of the Δtrl deletion strain compared to the isogenic rifampicin resistant *P. laumondii* DJC (DJCRif) at 28 °C and 36 °C.

### 2.2 The *trlG* gene is part of a small putative three gene operon

Examination of the genomic context of *trlG* revealed it is part of a small operon with *PluDJC_RS01880* and *PluDJC_RS01885*, encoded between the DNA polymerase gene, *polA*, and *dsbA* (Figure 1A). *PluDJC_RS01880* encodes a protein annotated as a histidinol phosphatase (WP_011144772.1). The product of *PluDJC_RS01885* (WP_011144773.1) on the other hand is a small 79 amino acid protein annotated as a hypothetical protein but homologues in other organisms are annotated as transcription factors.

To further confirm the role of *trlG* in temperature restriction we constructed an unmarked deletion mutant of the entire TRL operon, Δ*trl*. This mutant was also able to grow on both solid and liquid media at 36 °C (Figure 2B). Importantly, plasmid trans-complementation with the whole operon restored the temperature restricted phenotype in the SNP mutants (Figure 3). Note, we used the whole operon rather than just *trlG* alone to ensure the stoichiometry of the three gene members was preserved.

**Figure 3.**
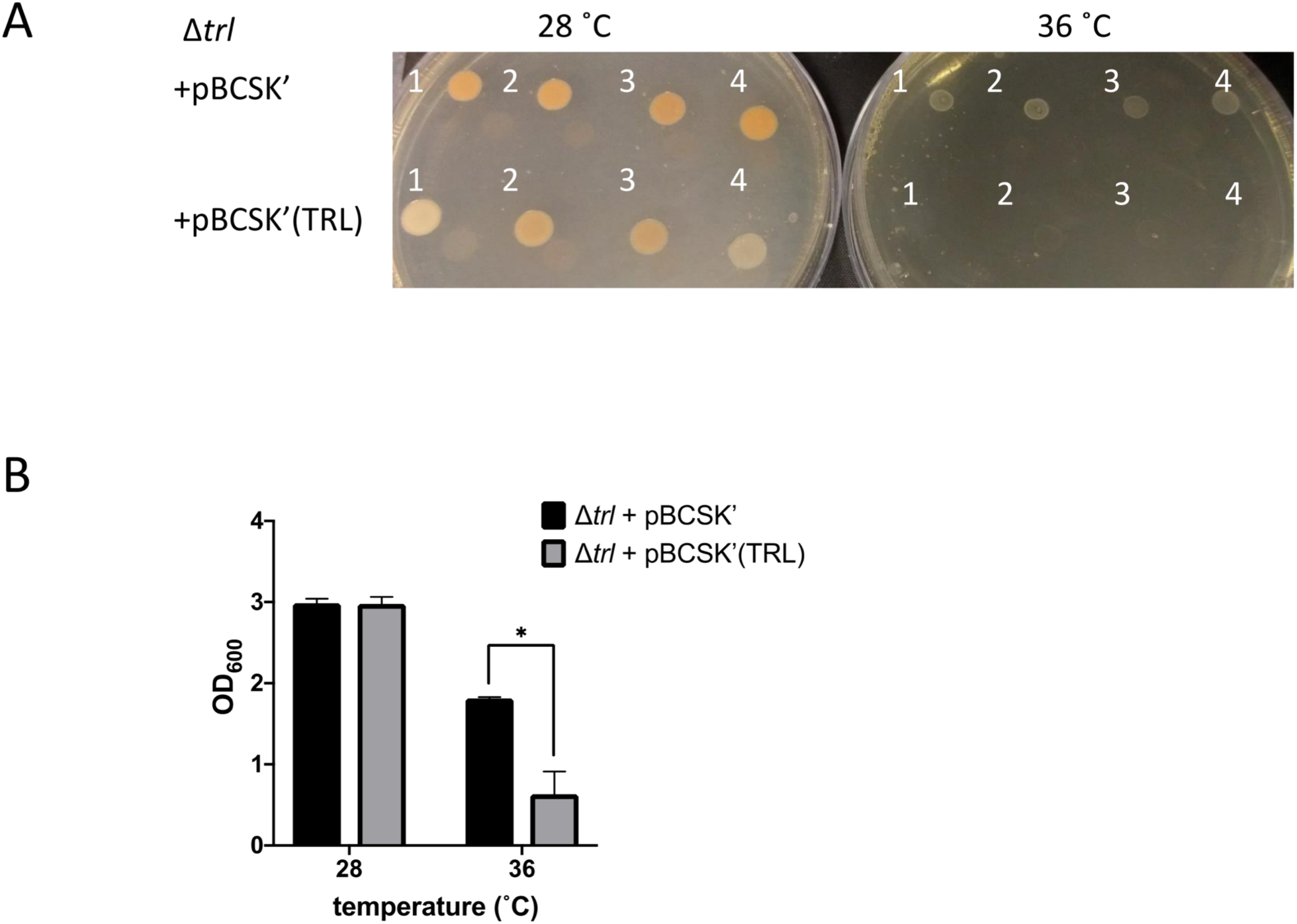
Growth of the Δ*trl* deletion mutant complemented with either the empty pBCSK’ vector or pBCSK’ carrying the TRL operon on A) LB agar and B) liquid LB media at 28 °C and 36 °C. The asterisk denotes significant difference at *p*_adj_ < 0.05. Results show the averages and standard error from four biological replicates.

Finally, since virulence of the bacteria to insects is a crucial aspect of their life cycle, we determined if mutation of *trlG* would have any effect on virulence against *Galleria mellonella* larvae. The experiment was performed at room temperature (∼20 °C), using a representative SNP mutant as well as the Δ*trl* deletion strain, compared to WT and the Rif_R_ isogenic parent strains respectively. Interestingly, we observed that the SNP mutant killed the *Galleria* larvae much faster than the WT (Figure 4). This was also the case for the Δ*trl*, albeit the effect was less pronounced. We also performed this virulence assay at 36 °C. As expected the WT strain was not able to kill the *Galleria*, although surprisingly, even though they could grow at this temperature *in vitro*, the mutant strains also failed to establish an infection.

**Figure 4.**
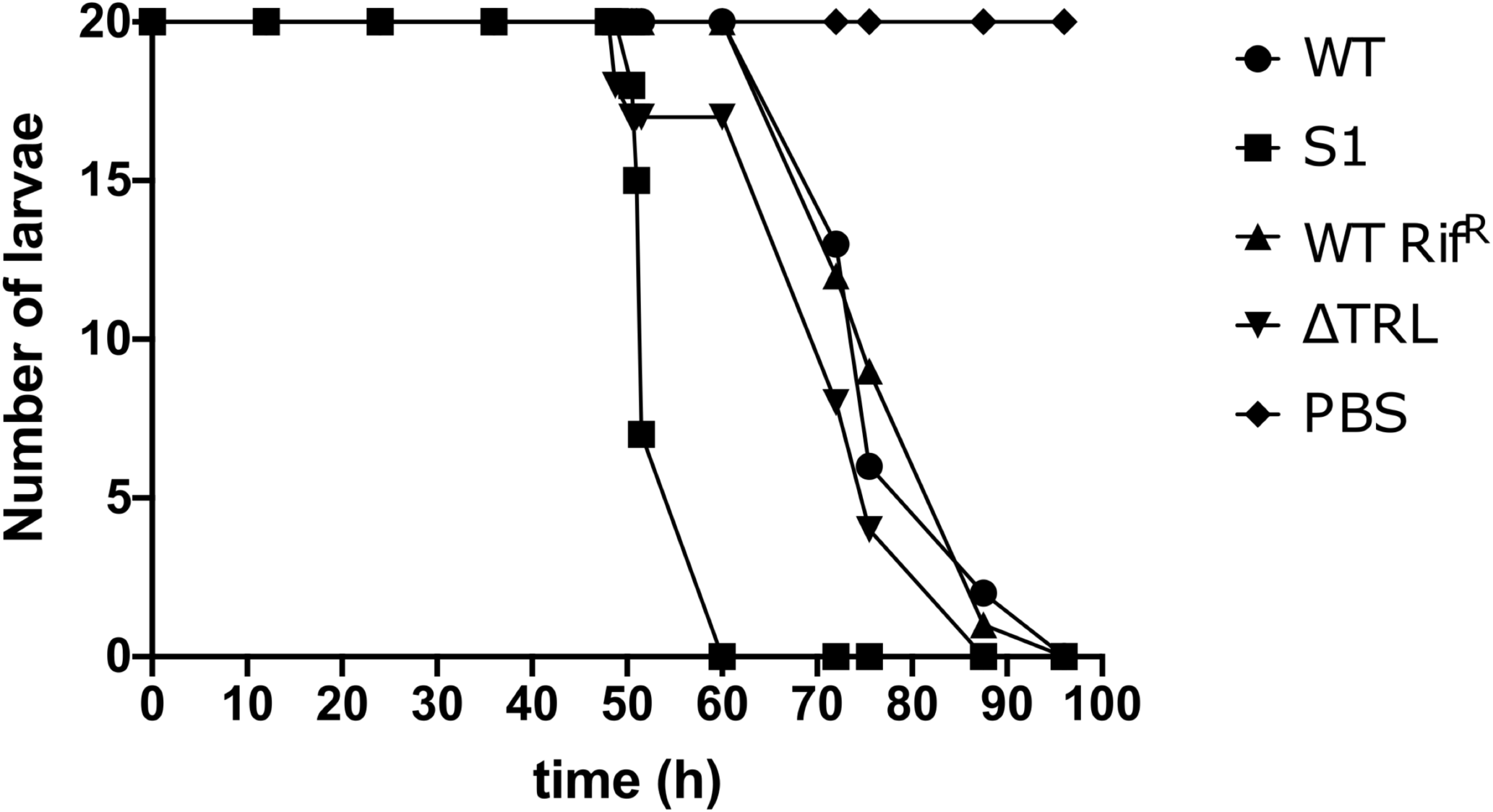
Virulence of the *trl* mutants. The results show the number of surviving *Galleria mellonella* larvae infected with either the WT strain, the S1 SNP mutant, the RifR WT strain or the Δtrl deletion mutant at different time points. *G. mellonella* larvae injected with PBS were used as control. Twenty larvae were used for each strain and the experiment was repeated with similar results.

### 2.3 Transcription of the TRL operon is induced by temperature up-shift

To determine whether the TRL operon is specifically expressed in response to exposure to the higher temperature we created reporter constructs whereby we cloned regions upstream of the operon upstream of a promoterless GFP gene. An *in silico* search for promoters using BPROM (Solovyev and Salamov 2011) showed the presence of two putative promoters upstream of the first gene in the TRL operon (*PluDJC_*RS01880, designated here as *trlR*). Interestingly, one of these promoters seems to be upstream of the predicted, oppositely encoded *polA* promoter. Analysis of apparent transcription start sites in our unpublished RNAseq analysis of *P. laumondii* supports the location of these predicted promoters. Additionally, a putative promoter is predicted which is encoded inside the *trlR* open reading frame. With TrlR itself showing homology to a transcriptional regulator we hypothesised that this might be a legitimate promoter driving expression of *trlF* and *trlG*. We therefore decided to create multiple constructs to ensure that we capture the transcriptional elements that control expression of genes in the TRL operon (Figure 5A). These constructs were separately introduced back into a rifampicin resistant isolate of DJC and the effect of temperature up-shift tested. These experiments confirmed that transcription of the operon is increased upon temperature up-shift (Figure 5B and 5C). It should be noted that the growth of *P. laumondii* DJC following temperature up-shift does not immediately cease, but rather continues to increase briefly before reaching a plateau (Supplementary figure 3A). Moreover, at the end of the 24 h period and following a ∼20 h incubation at the higher temperature, a large proportion of cells remain viable as shown by live/dead staining (Supplementary figure 3B). Importantly, the expression of these reporter constructs when the bacteria are grown at 28 °C was also monitored and we observed that the operon is also expressed at the permissive growth temperature but expression shows a gradual increase as the bacteria go through late exponential and stationary phase (Figure 5B and 5C).

**Figure 5.**
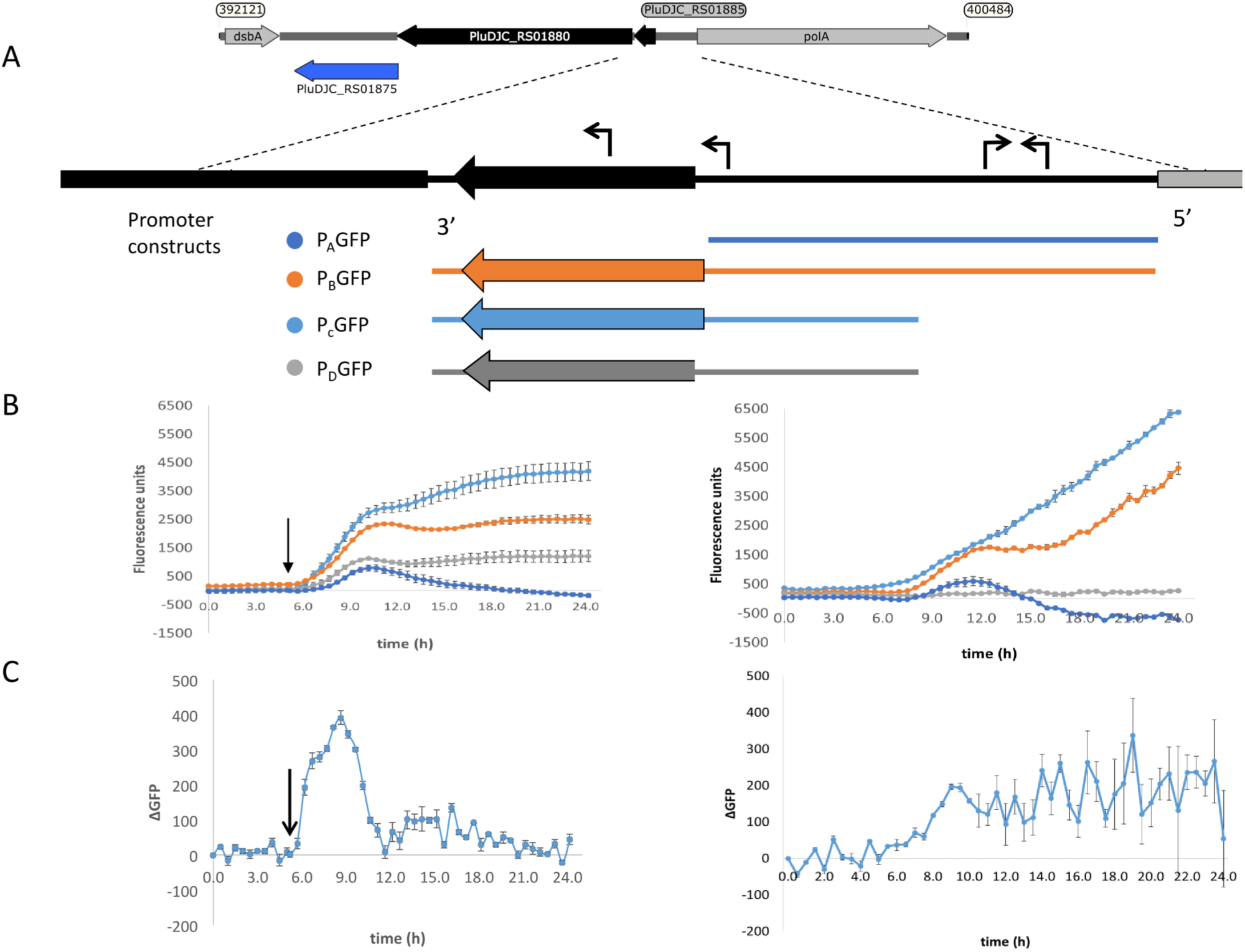
Up-regulation of the promoter of the TRL operon upon shift to 36 °C. A) Schematic diagram showing the different regions used in the promoter reporter constructs. The expanded panel shows the region between *trlF* (*PluDJC_RS01880*) and *polA* with the predicted promoters based on BPROM indicated as arrows on top of the sequence. At the bottom are the regions amplified and introduced separately into pGAG1 upstream of the *gfpmut3* gene to create promoter constructs P_A_GFP-P_D_GFP. The diagrams are color coded to reflect the colors of the data plotted below. B) Left: Fluorescence of the *P. laumondii* DJC reporter strains following a shift in temperature from 28 °C to 36 °C after 5 h of growth as indicated by the arrow. Right: Fluorescence of the *P. laumondii* DJC reporter strains grown continuously at 28 °C. Results are arbitrary fluorescence units following a subtraction of the fluorescence of the *P. laumondii* DJC carrying pGAG1 in the absence of any promoter to account for any autofluorescence and they represent the mean of three biological replicates +/- standard error. C)The rate of change in fluorescence in *P. laumondii* DJC carrying pGAG1(PCGFP), calculated by subtracting the fluorescence at a given time point by the fluorescence of the previous time point; left: following a shift in temperature from 28 °C to 36 °C after 5 h of growth as indicated by the arrow; right: at 28 °C. Results represent the mean of three biological replicates +/- standard error.

### 2.4 Presence of the TRL in the *Photorhabdus* genus

Since mutations in *trlG* appear to allow *P. laumondii* to survive in higher temperatures we were interested in investigating the presence of the gene among other *Photorhabdus* genomes. A *trlG* homologue was found in 21 out of the 46 *Photorhabdus* genomes in the RefSeq genome database (Figure 6 and Supplementary Figure 4). However, no *trlG* homologues were present in any bacteria belonging to the *P. asymbiotica* or *P. temperata* subgroups. Within the group of bacteria previously classed as *P. luminescens* the presence of the gene is not uniform. Those isolates that possess the gene actually encode for the full operon. Even though the majority of those genomes are in the draft sequencing stage, we investigated the location of the operon in the genome and observed that it appears to be conserved in the region between *dsbA* and *polA*. Importantly, in the published genome sequence of the type strain *P. laumondii* (Duchaud et al. 2003) an insertion sequence (IS) element is found upstream of the *polA* gene suggesting the operon may have been horizontally acquired. However, we note that this IS element is absent in our WT *P. laumondii* DJC strain – which is a laboratory selected derivative of the original TT01 isolate (confirmed by PCR). Interestingly, Alien Hunter analysis (Vernikos and Parkhill 2006) shows no evidence of horizontal gene transfer of this three gene TRL locus, suggesting it is ancestral in this sub-lineage of *Photorhabdus*. In addition to *P. laumondii* TT01, only *P. laumondii* strain DSPV002N possesses an IS630-like element ISPlu10 family transposase directly upstream of the operon. Four other isolates, *P. laumondii* HP88, *Photorhabdus* sp. S7-51, *Photorhabdus* sp. S15-56 and *Photorhabdus* sp. S14-60, appear to possess a gene encoding for a 61 amino acid hypothetical protein (WP_133148736.1), upstream of the operon but in the opposite strand.

**Figure 6.**
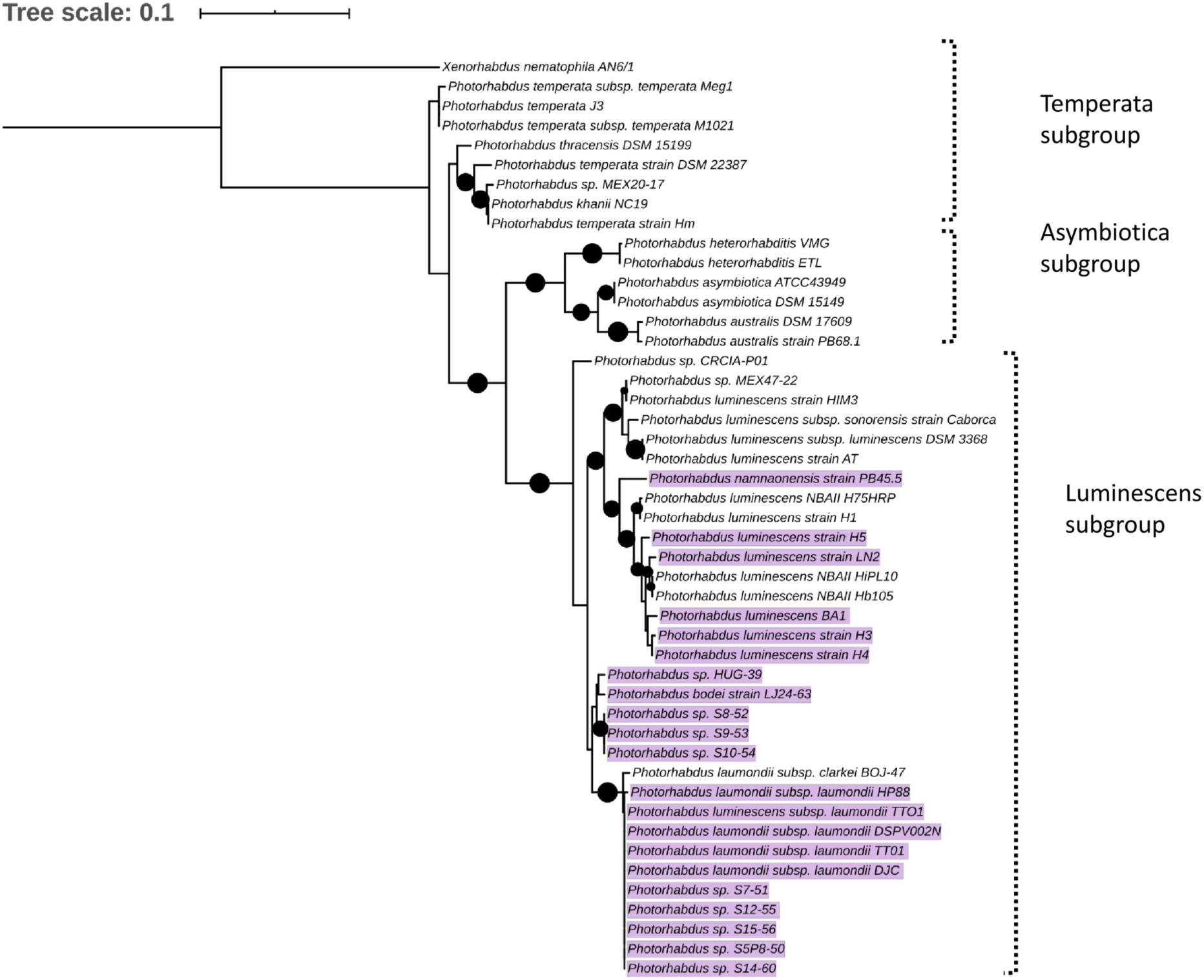
Maximum likelihood tree constructed using the *Photorhabdus recA* nucleotide sequences from RefSeq genomes. Black circles indicate branch support over 80 %. Highlighted are the strains that were found to contain a copy of the *trlG* gene.

### 2.5 The TRL is conserved amongst bacterial genera

The TRL is of relevance to bacteria beyond the *Photorhabdus* genus. Blastp analysis against the NCBI reference protein database showed that TrlG is prevalent in proteobacteria (Figure 7). The closest homologue outside of *Photorhabdus* is found in *Vibrio parahaemolyticus* followed by proteins encoded by *Pseudospirillum japonicum* and *Nitrincola nitratireducens*, which like *Photorhabdus* belong to the class of γ-proteobacteria. TrlG was, however, absent from the closely related *Xenorhabdus*, also a nematode endosymbiont. While the TrlG proteins of *Photorhabdus* and *Glaeserella* are all clustered together, for other bacteria this does not seem to be the case, suggesting individual acquisition events. Within the *Vibrio* genus, the gene is found in specific isolates of different species, such as *V. cholerae, V. anguillarum* and *V. atlanticus*. As an example, out of the 1218 *V. cholerae* genomes found in the RefSeq database, only 12 were found to encode for a homologue of TrlG (with over 40 % sequence identity). Of note is the presence of TrlG in various *Pseudomonas* isolates. Some are classified as strains of the plant pathogens *Pseudomonas syringiae* or *Pseudomonas amygdali*, however the gene is also present in some isolates of *P. aeruginosa*.

**Figure 7.**
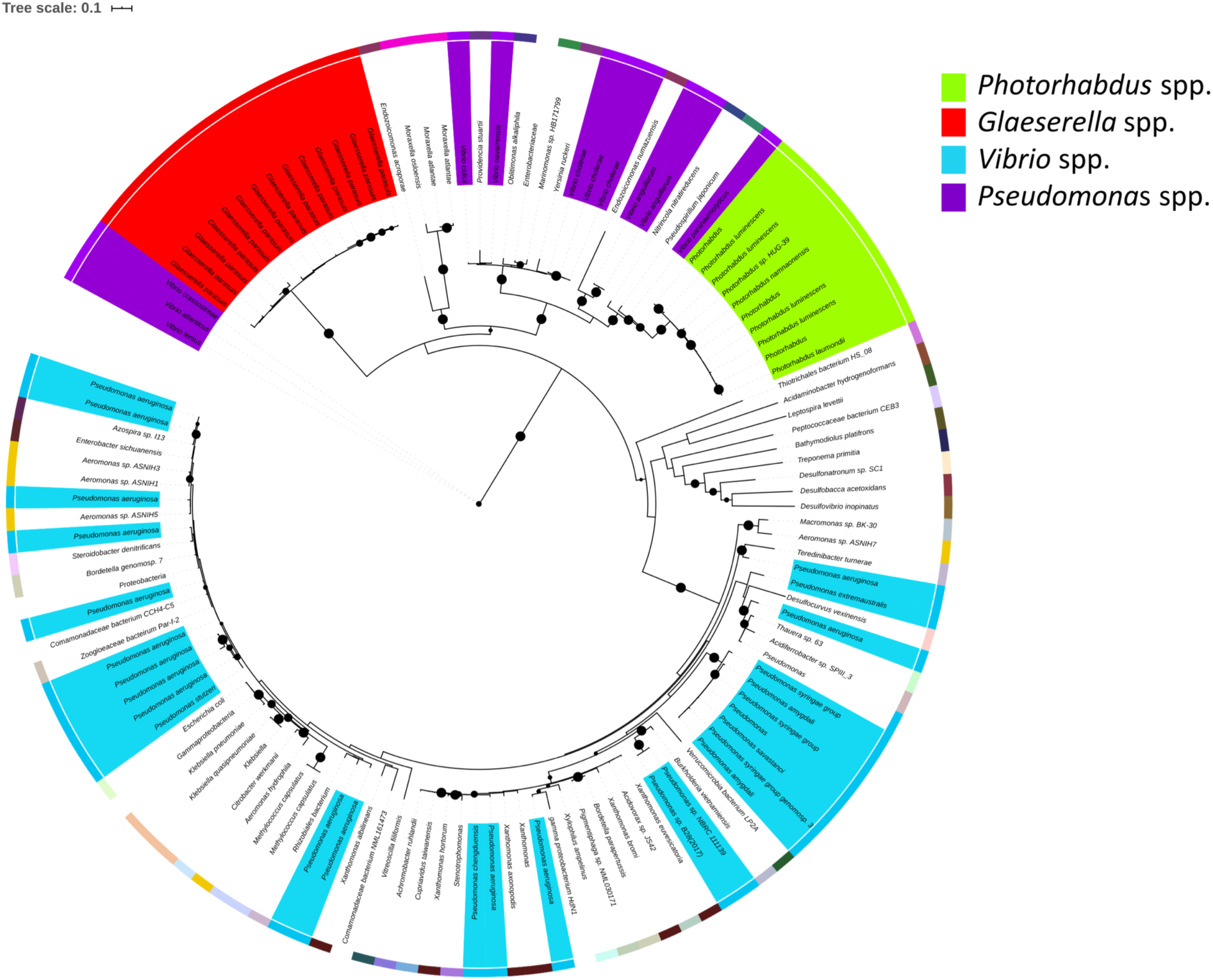
Maximum likelihood tree constructed using protein sequences with over 60 % sequence identity to the *Photorhabdus laumondii* TrlG protein (with a query cover > 90 %). Nodes with support values higher than 80 % are indicated with black circles. The external coloured ring represents different genera. Protein identifiers for all the entries can be found in Supplementary table 1.

Analysis of conservation of the entire operon using MultiGeneBlast (Medema, Takano, and Breitling 2013) against a Genbank database consisting of complete bacterial genomes showed that outside of the *Photorhabdus* genus, all three genes, *pluDJC_RS01875 - RS01885* are only conserved syntenically in strains of *Vibrio anguillarum* and on plasmid pYR3 of *Yersinia ruckeri* strain CSF007-82 (locus tags CSF007_p0360, CSF007_p0365, CSF007_p0345, Figure 8). However, unlike the TRL of *P. laumondii* where the three genes are located in succession, in pYR3 there are two small genes annotated as encoding Type I restriction-modification system, DNA-methyltransferase subunit M in between CSF007_p0345, the gene encoding the putative transcriptional regulator, and CSF007_p0360. *Yersinia ruckeri* is the causative agent of enteric red mouth disease in salmonid fish species (Kumar et al. 2015) and its optimum growth temperature is also 28 °C, while its optimum temperature for infectivity is 18 °C. Similarly, *V. anguillarum* is also a fish pathogen with an optimum growth temperature of 25 °C whilst the optimum temperature for vibriosis disease is 15 °C (Lages, Balado, and Lemos 2019). It appears that a lower temperature optimum for virulence is common in fish pathogens and virulence genes are found to be up-regulated at these lower temperatures (Guijarro et al. 2015; Mendez et al. 2018; Lages, Balado, and Lemos 2019).

**Figure 8.**
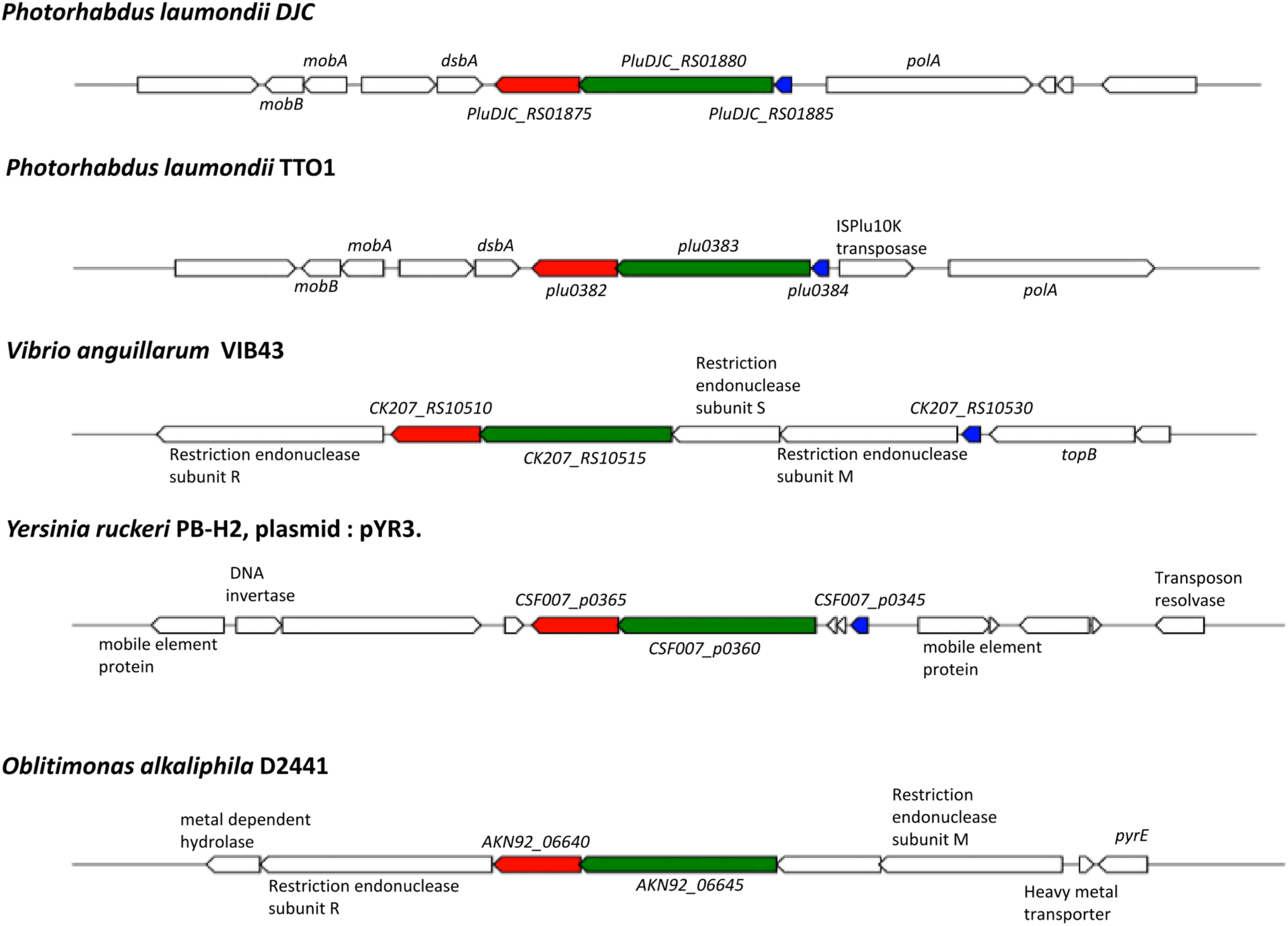
Conservation and genetic organization of the TRL operon in representative species as detected by Multigene blast. In red is *trlG*, in green is *trlF* and in blue is *trlR*; genes coloured identically in other organisms denote the identified homologues of *trlG, trlF* and *trlR* respectively.

## 3 Discussion

Members of the genus *Photorhabdus* dedicate a significant proportion of their genome to the synthesis of virulence factors and resource defence. Indeed, when the first *Photorhabdus* genome sequence was completed it was noted that there were more predicted toxin genes than in any other bacteria that had been sequenced at the time (Duchaud et al. 2003). In addition, many other genera belonging to the γ-proteobacteria contain multiple examples of mammalian pathogens, both obligate and opportunistic. With these observations in mind it is perhaps unusual that only very few members of the genus *Photorhabdus* are able to cause infection in humans (Gerrard and Stevens 2017). Our previous analysis of the adaptations relevant to human pathogenicity in a representative *P. asymbiotica* strain suggested that no specific mammalian active toxins are necessary to facilitate human infection. Rather it seems that substantive changes in metabolic activity, when cultured at 37 °C, are primarily responsible for their expanded host range (Mulley et al. 2015); a phenomenon termed nutritional virulence (Abu Kwaik and Bumann 2013).

An obvious limitation for mammalian infection in many members of the genus is their inability to grow above ∼35 °C, which again is unusual for members of the γ-proteobacteria. Typically, maximum growth temperatures of bacteria are determined by multiple factors that affect bacterial physiology, such as membrane and lipid fluidity, chaperone activities and other enzyme/protein functions. It is therefore possible that for any one species there are multiple functions that fail simultaneously at the non-permissive temperature. However, the ability of *P. asymbiotica* isolates to grow readily at temperatures of 37 °C (and above) and the overall high level of conservation in core genome sequence across the genus indicates that there is likely no general physiological reason for temperature restriction in members of the genus.

Our identification of the TRL genes offers up an alternative explanation to this temperature restriction phenomenon, at least in certain lineages of *P. laumondii*. The specific induction of the TRL gene transcription upon temperature up-shift, leading to growth arrest, appears *prima facia* to represent a specific adaptive mechanism. We do not yet know if induction of TRL transcription at higher temperatures is mediated by the TrlR protein alone or if other regulatory pathways are involved. While it remains unclear as to what advantage this temperature dependent growth arrest imparts upon these strains, we may speculate. For example, we note that the nematode symbiont cannot survive above ∼34 °C *in vitro*, although to our knowledge nematode survival at different temperatures *in vivo* has not been investigated. Therefore, a TRL mediated mechanism to suppress continued exponential growth at increasing temperatures, may represent a mechanism to synchronise life cycles of the bacteria and nematode in fluctuating environmental conditions. It is perhaps no coincidence that the TRL operon is encoded next to the *polA* gene and that the most 5’ TRL promoter overlaps on the opposing strand with that of the *polA* gene. Thus, activation of TRL transcription might directly suppress *polA* transcription by steric hindrance. It is formally possible that the tight linkage between *polA* and TRL reflects a functional linkage in the gene activities (ie, replication and temperature). It is interesting in this respect that even at 28 °C, we see a gradual increase in TRL transcription as the culture moves toward stationary phase and the eventual cessation of replication.

Interestingly, while ablation of the whole TRL operon did not strongly affect virulence in the *Galleria* infection model at 28 °C, the S1 SNP mutant actually became significantly more virulent at this temperature. This suggests a functional interplay between *trlG, trlF* and *trlR*, as only the *trlG* gene is mutated in S1. Therefore, the activity of *trlF* and/or *trlR* in the absence of the balancing effect of *trlG* seems to enhance virulence activity. Importantly, when the host temperature was increased to 36 °C, the infection failed of both the S1 SNP and Δ*trl* strains, despite their ability to grow *in vitro*. This is consistent with the reported failure of *P. asymbiotica* to establish a lethal infection in the *Manduca sexta* model when incubated at 37 °C (Mulley et al. 2015). In a natural context 37 °C is not a problem for *P. asymbiotica* ATCC43949 as it would be in a mammalian host, for which it is adapted. Thus, despite the alleviation of growth restriction, there are clearly other temperature dependent changes in gene regulation occurring, as evidenced by the down regulation of luciferase activity and pigment production in these mutants. This effect may be common to both *P. asymbiotica* and the mutant *P. laumondii* lineages and regulated through a separate unknown mechanism. In addition, the loss of pigmentation and reduction in bioluminescence of the mutant colonies at 36 °C is reminiscent of the secondary phenotypic variants of *Photorhabdus* (Boemare and Akhurst 1988) as well as Δ*mdh* and Δ*fumC* mutants deficient in enzymes of the TCA cycle (Lango and Clarke 2010) and a Δ*relAspoT* strain, deficient in (p)pGpp synthesis (Bager et al. 2016). The aforementioned mutants retain virulence in insect models but are no longer symbiotically competent. Absence of pigmentation and bioluminescence is also a characteristic of the M-phase variants of *P. luminescens*, which are responsible for transmission of the bacteria to infective juvenile nematodes and establishment of symbiosis, however those variants are avirulent against insects (Somvanshi et al. 2012).

The majority of random mutations selected from the population were non-synonymous amino acid substitutions in TrlG rather than premature stop codons or frame shifts. It is possible that this simply reflects the lower probability of obtaining a stop codon by random chance as complete deletion of the operon suggested the presence of an altered version of TrlG was not in some way advantageous. More importantly, no mutations were seen in the much larger and tightly linked *trlF* gene. This suggests that either (i) TrlF does not play a role in the temperature restriction phenotype, or (ii) that if it does, its function can be fully compensated by dominant activity of an intact TrlG protein. The small size of the *trlR* regulator gene means there was a much lower probability of isolating mutants in this. As we could delete the whole operon, it indicates that mutations in *trlF* alone are not overtly deleterious if they should occur in isolation.

As the lifestyles of most *Photorhabdus* strains are generally similar, it is interesting that the TRL operon appears restricted yet ancestral to a specific species lineage of *Photorhabdus*. As members of this linage are geographically dispersed (Machado et al. 2018; Mathur et al. 2018), factors such as biogeography, insect prey type or the nature of competing organisms are unlikely to be the reason for maintaining TRL. We therefore speculate that selection for TRL may be a result of specific requirements of the host bacterial genomes, or perhaps in restrictions imposed by the nematode host lineage. For example; as the *P. luminescens* TT01 genome is heavily infested by IS-elements, we could hypothesise that TRL represents a mechanism to protect against rampant IS-element activity. If a sustained temperature increase was “perceived” as stress to the bacteria (or more importantly the IS-elements), it may lead to undesirable activation of a large number of IS-elements. Therefore, stopping genome replication would likely protect against extensive transposition damage.

An alternative idea is that these strains rely on the activities of a pathway or global regulator which is highly sensitive to temperature. For example, it has been shown that secondary metabolite production is reliant upon the activity of the Hfq protein (Tobias et al. 2017) which, in other Gram-negative bacteria studied, mediates interactions of regulatory ncRNAs with their target mRNA molecules (A. Zhang et al. 1998; Møller et al. 2002; Lease and Woodson 2004). Folding and interactions of many ncRNAs are often temperature sensitive. The lack of pigment and bioluminescence in the mutant strains at the higher temperature suggests that this regulation is indeed being perturbed. If all secondary metabolism necessary for virulence and symbiosis ceases at the higher temperature it would make sense for the bacterium to inhibit further replication until normal regulation is re-established.

Finally, the identification of TRL-like operons in members of other genera suggests this represents a widespread general mechanism, regulating the life-cycles of phylogenetically diverse bacteria. The association of TRL operons with pathogens of poikilothermic hosts (e.g. fish) tempts speculation there is an as yet unknown advantage to this mechanism in such niches.

## 4 Materials and Methods

### 4.1 Strains and growth conditions

The strains used in this study were the rifampicin sensitive parent strain of *P. laumondii* subsp. *laumondii* DJC (WT) (Zamora-Lagos et al. 2018), and a rifampicin resistant derivative of this strain, *P. luminescens* spbsp. *laumondii* DJC (Rif_R_) (Jones et al. 2010). Bacteria were grown in Lysogeny broth (LB) at 28 °C unless otherwise stated. For growth on solid media, LB was supplemented with 1.5 % agar and 0.1 % sodium pyruvate and plates were incubated in the dark. When necessary, kanamycin was added to the media at a concentration of 25 µg/ml. To assess growth on solid media, overnight cultures of *Photorhabdus* were diluted to an OD_600_ of 0.05 and 5 μl were spotted on solid media. They were then incubated at the appropriate temperature for 48 h. To assess growth in liquid, the diluted cultures were incubated at the appropriate temperature for 24 h, shaking at 180 rpm.

### 4.2 Isolation of the temperature tolerant clones

Saturated overnight cultures of *P. laumondii* DJC grown at 28 °C in LB medium were plated on LB agar (LBA) and incubated at 36 °C. After approximately 48 h colonies were observed on the plates. These were re-streaked 4 times on LBA at 36 °C and stably grown clones were stored at -80 °C in 32 % v/v glycerol.

### 4.3 Whole-genome sequencing and bioinformatics analysis

DNA was isolated from 1 ml of overnight cultures of *P. laumondii* DJC using the Qiagen Blood and Tissue kit. The Nextera XT kit (Illumina) was used for library preparation and sequencing was performed using a MiSeq v2 cartridge on a MiSeq sequencer (Illumina). Raw data has been deposited in the NCBI database under BioProject ID PRJNA596169, while raw sequencing reads for the WT isolate in our lab can be found in the short read archive using accession number SRR8307122. Fastq reads of both the WT and the tolerant clones were mapped against the NCBI reference sequence for *P. luminescence* subsp *laumondii* DJC NZ_CP024900.1(Langmead and Salzberg 2012) in very sensitive mode. SAMtools v0.1.18 (Li et al. 2009) was used to obtain sorted bam files, containing aligned reads, and subsequently pile up format files. VarScan v2.3.6 (Koboldt et al. 2012) was then used to identify SNPs (parameters: p-value 0.1, min-coverage 20 --min-reads2 15 --min-avg-qual 30 --min-var-freq 0.9). A different set of parameters was subsequently used to account for the clones with lower coverage (with modified min-coverage of 10 and --min-reads2 of 6). For clones where no SNP was identified in the *trlG* gene automatically, manual inspection of the alignment files followed by PCR and Sanger sequencing were used to check for the presence of any insertions/deletions. In particular, to sequence the *trlG* gene in the JC and JD strains as well as the tolerant clones S15 and S29, either the gene was amplified using primer pair 00323SecF/00323SecR or the entire operon was amplified using recOSeqF/reOSeqR and the resulting PCR products were purified and sent for Sanger sequencing. All primer sequences used in this study are listed in Table 2.

**Table 2:**
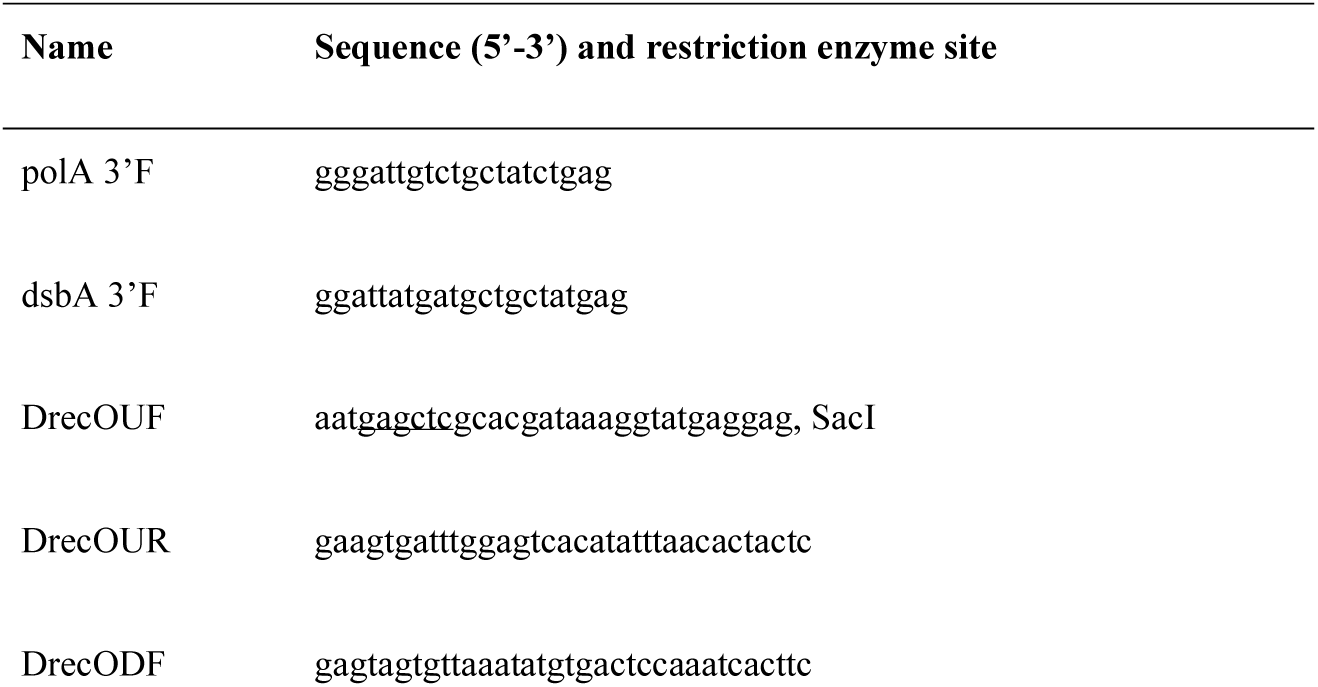
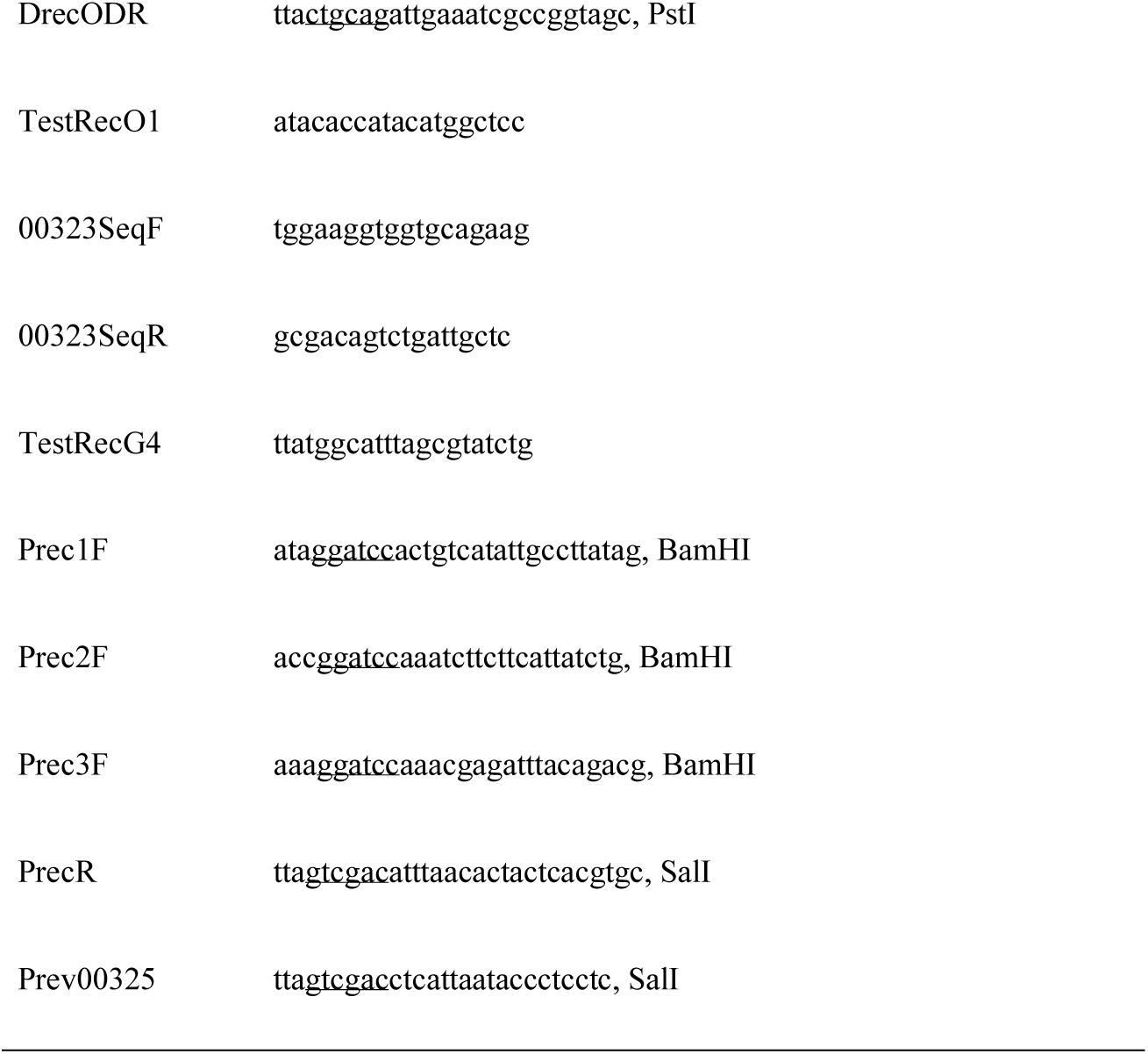
Oligonucleotide primers used in this study. The positions of restriction sites are shown underlined.

A prediction of the protein structure of TrlG was performed using I-TASSER (Y. Zhang 2008) with default parameters. The location of the amino acid substitutions on the protein was visualised using UCSF Chimera (Pettersen et al. 2004).

To investigate the presence of the gene in other bacteria, *trlG* was translated using the standard codon usage and the amino acid sequence was used as input for BLASTP v2.8.1 against the NCBI protein reference sequences using default search parameters. Hits were retained if the showed at least 60 % sequence identity over 90 % of the query length. Multiple alignments were created using Clustal Omega (Sievers et al. 2011). The Clustal alignment file was then used to infer a maximum likelihood phylogeny using IQ-TREE v 1.6.3 (Nguyen et al. 2015) with the JTT substitution model selected by ModelFinder (Kalyaanamoorthy et al. 2017), and ultrafast bootstrap (Hoang et al. 2018). To reconstruct the *Photorhabdus recA* phylogeny, *recA* nucleotide sequences were obtained from the *Photorhabdus* genomes present in the NCBI reference genomic sequence database and analysed using the Phylogeny.fr platform. The sequences were aligned using MUSCLE and ambiguity was removed using Gblocks. A maximum likelihood tree was then constructed using PhyML and 500 bootstrap iterations were performed. Phylogenetic trees were visualised using the Interactive Tree Of Life (iTOL) (Letunic and Bork 2019). For MultiGeneBlast analysis (Medema, Takano, and Breitling 2013), a database was created from the bacterial Genbank division and a homology search was performed with default parameters, with the three TRL genes as input.

### 4.4 Construction of a Δ*trl* mutant

A markerless Δ*trl* deletion mutant was created by double homologous recombination using the suicide plasmid pDS132 (Philippe et al. 2004). Regions upstream and downstream of the TRL were amplified using primer pairs DrecOUF/DrecOUR and DrecODF/DrecODR respectively. The purified PCR products were mixed and used as a template in a second joining PCR with primers DrecOUF/DrecODR. The resulting product was digested using enzymes SacI and PstI and ligated into digested and dephosphorylated pDS132. The resulting plasmid was introduced by transformation into the donor *E. coli* strain S17.1 lamda-pir. For conjugations into *Photorhabdus*, the rifampicin resistant *P. luminescens* DJC strain was used as a recipient. Briefly, exponentially growing cultures of *P. laumondii* DJC Rif_R_ in LB containing 1 mM MgCl_2_ and *E. coli* S17.1 lamda-pir harbouring pDS132(Δ*trl*) were harvested and combined at a ratio 4:1 of acceptor to donor bacteria. Matings were spotted on LBA plates containing 1 mM MgCl_2_ and incubated overnight at 28 °C. Bacteria were then harvested and aliquots were plated on LBA supplemented with 15 µg/ml chloramphenicol and 50 µg/ml rifampicin. Possible transconjugants and hence first recombinants were confirmed by PCR using primer pairs TestRecO1/00323SeqR and polA3F’/ TestRecG4. Successful first recombinants were grown in LB overnight and then the culture was diluted in fresh LB and grown to mid exponential. At this point serial dilutions were made and aliquots were plated on LBA supplemented with 0.2 % sucrose and incubated at 28 °C. The resulting colonies were tested for deletion of the TRL operon by PCR using primers TestRecO1/TestRecG4.

### 4.5 Complementation

For complementation, the entire TRL was amplified using primers recOFEcoRI/recORSpeI and the resulting PCR product was digested with EcoRI and SpeI and ligated into a modified version of vector pBCSK (that had been previously digested with AseI and KpnI to remove Plac). The resulting plasmid, termed pBCSK’(TRL), and the empty pBCSK’ were introduced separately into the Δ*trl* deletion mutant or into SNP mutants by transformation. To prepare cells for transformation, bacteria growing exponentially in 100 ml LB were harvested at an OD_600_ of 0.2-0.3 and kept for 90 min on ice. They were then collected by centrifugation and washed twice with ice-cold SH buffer at pH 7 (10 mM HEPES containing 5 % w/v sucrose). The cells were resuspended in 160 µl SH buffer and 50 µl aliquots were used for electroporation with the following conditions: 2.5 kV, 25 μF and 200 Ω in 2 mm cuvettes. This was followed by addition of 1 ml LB and incubation at 28°C, 180 rpm for 3 h. Aliquots were plated on LB agar supplemented with 25 µg/ml chloramphenicol. Colonies were tested for the presence of the plasmid by PCR using primers M13R/Seq03223F.

### 4.6 Infection assays

The infections were performed using 5^th^ instar larvae of *Galleria mellonella*. The bacteria were grown in LB at 28 °C and then collected and resuspended in PBS to an OD600 of 1.5. The suspension was then serially diluted to obtain a final suspension with an infection dose of ∼50 colony forming units per 10 μl. The *Galleria* larvae were infected by injection through the first proleg and incubated either at room temperature or 36 °C. For the experiment conducted at room temperature two groups of ten larvae were infected per strain, while for the experiment at 36 °C, 8 larvae were infected per strain. The experiment was conducted twice.

### 4.7 Construction of GFP promoter fusions

The TRL promoter regions were amplified using Q5 DNA polymerase and with *P. laumondii* DJC DNA as a template and using the following primer pairs: Prec2F/PrecR for promoter region PA, Prec2F/Prev00325 for promoter region PB, Prec1f/Prev00325 for promoter region PC and Prec3F/Prev00325 for promoter region PD. PCR products were digested using BamHI and SalI and purified. They were then ligated into the broad range vector pGA-G1, which carries a promoterless *gfp*-mut3 (Leo Eberl, University of Zurich). The ligations were introduced by transformation into *E. coli* DH5a lamda-pir. Successful transformants were confirmed by PCR and plasmids were isolated using the Qiagen miniprep kit and introduced into *E. coli* strain S17.1 lamda pir to be used as a donor in conjugation, which was performed as described above.

### 4.8 Growth and fluorescence measurements of reporter strains

*P. laumondii* DJC strains harbouring the reporter constructs were grown overnight in LB supplemented with kanamycin at 28 °C and the cultures were then diluted to fresh media to an OD600 of ∼0.05. The diluted cultures were aliquoted in black flat-bottomed µClear 96-well plates (CELLSTAR, Greiner) and placed in an Omega Fluostar (BMG Labtech) microplate reader. They were incubated at 28 °C with double orbital shaking at 300 rpm. Readings of absorbance at 600 nm as well as GFP fluorescence (gain 1000) were taken every 30 min. For the temperature shift experiment, after 5 h of growth at 28 °C the temperature of the microplate reader was increased to 36 °C (this took place in ∼ 2 min). Then readings were taken immediately following the shift and at 30 min intervals after that.

## 5 Acknowledgements

We would like to thanks Prof. Mark Pallen for the use of his MiSeq machine.

## 7 Author Contributions

AH and NW designed the study; AH, JH and GM performed the mutant isolation experiments; JH performed the structural prediction studies; AH performed the remaining experiments and analysed the data. AH and NW wrote the initial manuscript draft. GM reviewed the manuscript. All authors have approved the final manuscript.

## 11 Supplementary material

Supplementary Figures 1-4

Supplementary Table 1

## 12 Data Availability Statement

The raw sequencing data generated and analysed in this study has been deposited in the NCBI database under BioProject ID PRJNA596169, while raw sequencing reads for the WT isolate in our lab can be found in the short-read archive (SRA) using accession number SRR8307122. The reference genome sequence for *P. luminescence* subsp *laumondii* DJC is available on NCBI RefSeq with the accession number NZ_CP024900.1.

