## Supplementary figure 1 for "Temperature restriction in entomopathogenic bacteria"

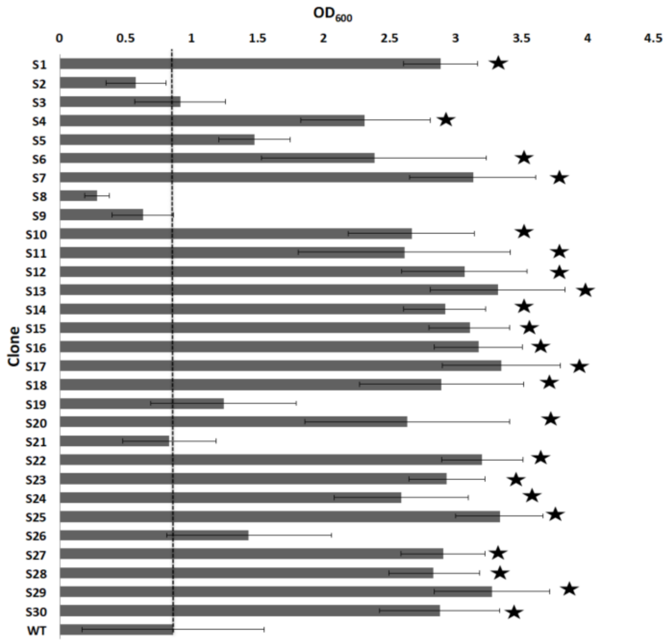


**Supplementary Figure 1**. Growth of the isolated clones in liquid at 36 ˚C after 24 h. The results show the averages of four independent replicates +/- standard error. The clones marked with stars are the ones in which *trlG* was confirmed to possess a SNP or be otherwise disrupted.
