## Supplementary figure 2 for "Temperature restriction in entomopathogenic bacteria"

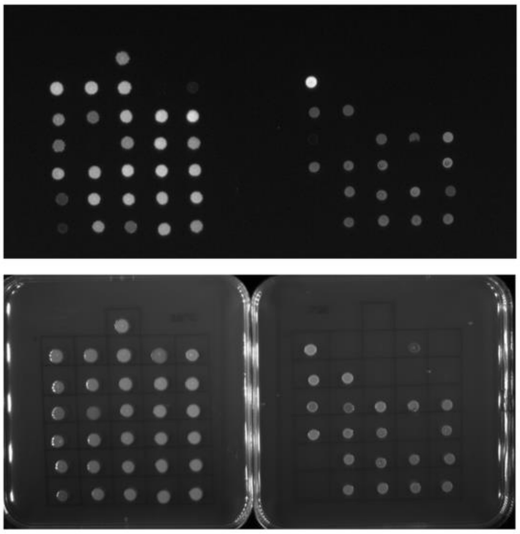

**Supplementary Figure 2.** Luminescence of the WT and the 30 sequenced tolerant clones shown in Figure 4A at 28 ˚C and 36 ˚C imaged using the GENESys SYNGENE imager (Top). At the bottom is the corresponding image of the plates taken with white light as a reference.
