## Supplementary figure 3 for "Temperature restriction in entomopathogenic bacteria"

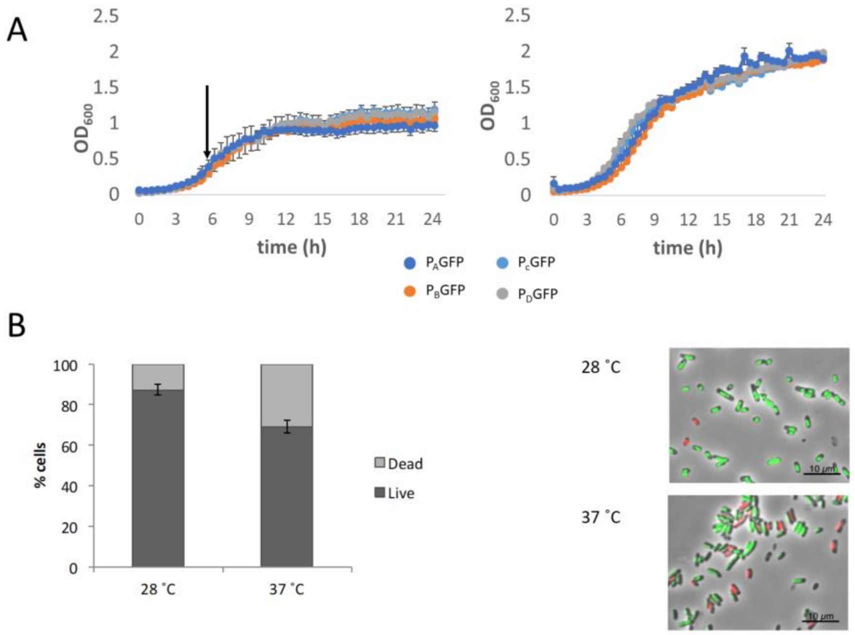


**Supplementary Figure 3**. A) Growth of *P. laumondii* DJC reporter strains following shift to 36 ˚C at the time indicated by the arrow (left) or at 28 ˚C (right) B) Proportion of live and dead cells of *P. laumondii* DJC WT after shift to 37 ˚C and incubation at the higher temperature for 20 h as determined by Baclight Live/Dead staining. The data are from two different samples for each temperature with 4 images per sample with at least 100 bacteria per image. Counting was done automatically using the Fiji software on colour thresholded images and using particle analysis with the size set between 0.01-0.2.
