## Supplementary figures and images for "Temperature restriction in entomopathogenic bacteria"

### Supplementary figure 4

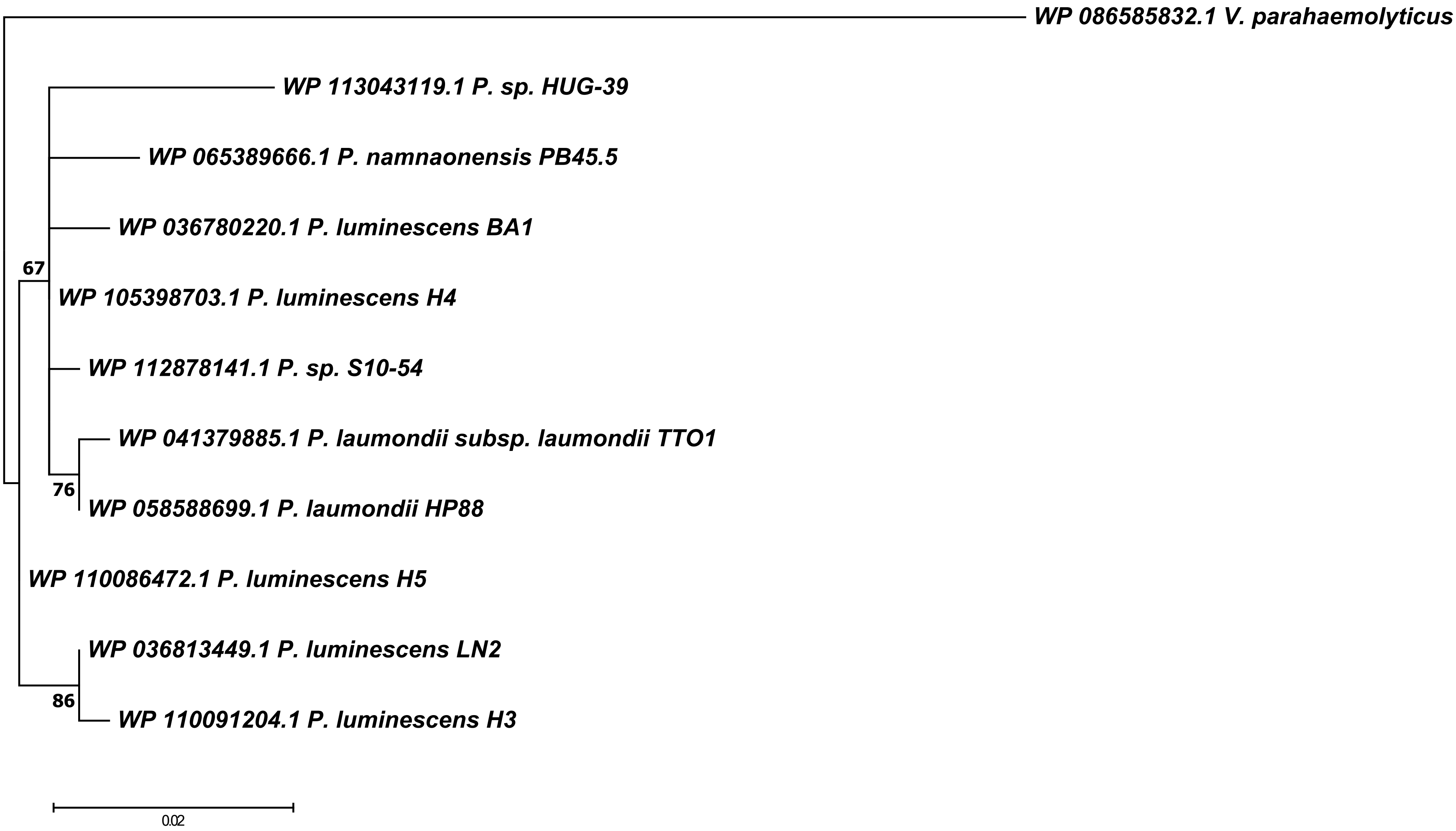
